# An ecological investigation of the effect of background noise on speech processing in a Virtual Classroom

**DOI:** 10.1101/2024.01.05.574300

**Authors:** Orel Levy, Adi Korisky, Yair Zvilichovsky, Elana Zion Golumbic

## Abstract

Many real-life situations can be extremely noisy. Psychoacoustic studies have shown that background noise can have a detrimental effect on the ability to process and understand speech. However, most studies use stimuli and task designs that are highly artificial, limiting their generalization to more realistic contexts. Moreover, to date, we do not fully understand the neurophysiological consequences of trying to pay attention to speech in a noisy place.

To address this lab to real-life gap and increase the ecological validity of speech in noise research, here we introduce a novel audiovisual Virtual Reality (VR) experimental platform. Combined with neurophysiological measurements of neural activity (EEG), eye-gaze and skin conductance (GSR) we studied the effects of background noise in a realistic context where the ability to process and understand continuous speech is especially important: A VR Classroom.

Participants (n=32) sat in a VR Classroom and were told to pay attention to mini-lecture segments by a virtual teacher. Trials were either Quiet or contained background construction noise, emitted from outside the classroom window, which was either Continuous (drilling) or Intermittent (air hammers).

Result show that background noise had a detrimental effect on learning outcomes, which was also accompanied by reduced neural tracking of the teacher’s speech. Comparison of the two noise types showed that the intermittent construction noise was more disruptive than continuous noise, as index by both behavioral and neural measures, and it also elicited higher skin-conductance levels, reflecting heightened arousal. Interesting, eye-gaze dynamics were not affected by the presence of noise.

This study advances our understanding of the neurophysiological effects of background noise and extends it to more ecologically relevant contexts. It also emphasizes the role that temporal dynamics play for processing speech in noise, highlighting the need to consider the features of realistic noises, as we expand speech in noise research to increasingly realistic circumstances.

## Introduction

The primary modality of learning in frontal classroom settings requires listening and paying attention to the teacher’s speech for long periods of time (Martin & Miller, 2012). However, school classrooms, like many other real-life environments, can sometimes be extremely noisy, with sounds from the outside penetrating through the windows and walls and interfering with ongoing class activities (Sala & Rantala, 2016; Shield et al., 2015; Shield & Dockrell, 2004). For instance, a study done in 2016 showed that the average noise level during classroom instruction in schools was 55 db at least half of the time (Sala & Rantala, 2016). The challenge of learning in noisy places is not limited to classrooms, but extends to remote learning as well (such as learning over Zoom, as experienced by many during the recent COVID-19 crisis), where students cannot always control the acoustic environment around them. Needless to say, noise can be extremely detrimental to learning, as noise severely impairs speech intelligibility, makes it more difficult to maintain attention to the lesson and ultimately has been shown to reduce academic performance (Astolfi et al., 2019; Klatte et al., 2010; Sala & Rantala, 2016).

Behavioral studies have shown that even moderate level of noise can substantially reduce speech intelligibility, comprehension, communication, and learning outcomes (Darwin, 2008; Dubbelboer & Houtgast, 2007; Sala & Rantala, 2016). These behavioral effects are also mirrored in less effective neural processing of speech in noise (Kong et al., 2015; Vanthornhout et al., 2018; Zou et al., 2019). Many studies have shown that attempting to understand speech in noisy environments is often accompanied by neural and physiological responses associated with heightened listening and attentional effort, such as increased neural alpha-band power (Becker et al., 2013; Obleser et al., 2012; Strauß et al., 2014; Wöstmann et al., 2015), increased skin-conductance responses (Mackersie & Cones, 2011), changes in the pupil diameter (Koelewijn et al., 2012; Miles et al., 2017) and the pattern of eye-gaze dynamics (Król, 2018; Šabić et al., 2020).

However, most research on the effects of noise on speech processing uses highly artificial task designs (e.g., simple word recognition), simplistic stimuli (e.g., word lists or short sentences) and impoverished sensory contexts (e.g., auditory-only presentation while looking at a blank computer screen). These do not fully capture the sensory and cognitive challenges of real-life environments, which are inherently multisensory, dynamic and highly contextual and therefore limited in their ecological validity (Holleman et al., 2020). In attempt to enhance our understanding of how the brain deals with speech in realistic noisy environments, and specifically during learning, here we use a Virtual Reality (VR) Classroom experimental design to simulates a realistic learning context and study the effects of different types of background noise on neural processing, physiological responses and comprehension of the lesson (Adams et al., 2009; Brown et al., 2023; Liu et al., 2021; Tromp et al., 2018).

We focused on a genre of background noise that is extremely prevalent in many real-life environments, and is completely out of the listener’s realm of control: construction noise. We examined the effects of two types of construction noise that differ in their temporal characteristics - continuous noise from drilling, which is relatively constant over time, and intermittent noise of air-hammers, which consists of a sequence of pseudo-rhythmic bursts of noise. This choice was motivated by **two competing hypotheses** regarding how the temporal structure of noise may affect the severity of its interference with speech processing.

The **‘glimpsing hypothesis’** for speech-in-noise processing suggests that when background noise is intermittent and contains short ‘noiseless gaps’, this allows listeners to listen in the gaps and fill in the missing speech information masked by the noise (Cooke, 2006; Fogerty et al., 2018; Howard-Jones & Rosen, 1993). Accordingly, speech comprehension is expected to be better on the background of intermitted noise, relative to continuous noise where there is constant masking of the speech stimulus. However, an alternative **‘habituation hypothesis’** posits that it is easier to process speech on the background of continuous noise, as its monotonic nature leads to habituation over time, which ultimately makes continuous noise less disruptive (Banbury & Berry, 1997; Bell et al., 2012; Thompson & Spencer, 1966). This is in contrast to intermittent noise, with its frequent onsets and offsets of noise, that trigger repeated phase-resets of cortical and arousal responses (Doelling et al., 2019; Ten Oever et al., 2017), ultimately reducing cortical adaptation and leading to greater disruption of speech processing.

To date, few and conflicting findings have been reported when directly comparing the effects of continuous and intermittent noise on speech processing, although both types have been shown to have detrimental effects on speech processing relative to no-noise conditions (Eqlimi et al., 2020; Hygge et al., 2003; Kozou et al., 2005; Niemczak & Vander Werff, 2019). While some have found better intelligibility for speech on the background of intermittent vs. continuous noise (Koelewijn et al., 2012; Miller & Licklider, 1950), others have found that intermittent noise impaired performance and reduced the neural response to speech, relative to continuous noise (Maamor & Billings, 2017; Niemczak & Vander Werff, 2019). However, some of the discrepancies between these studies may be due to the specific parameters of the noise-stimuli used (mostly amplitude modulated white noise). In the current study, we chose to use recordings from real-life construction sites as our continuous and intermittent noise-stimuli, rather than simpler simulated noises, as a means to enhance the ecological validity and relevance of our research for real life settings.

Participants were immersed in a VR Classroom environment, where they sat in the center of the classroom surrounded by other classmates and listened to content vignettes on topics related to history and science from an animated avatar-teacher standing in front of the class (∼40 second long trials; Figure 1). After each trial, they answered comprehension questions regarding the content of the teacher’s speech. Trials could either be quiet or be accompanied by continuous or intermittent construction noise, emitted from a location outside the classroom. We collected a variety of metrics as participants engaged in learning in the VR classroom, including their neural activity (using electroencephalography; EEG), eye-gaze (using an eye-tracker embedded in the VR headset) and galvanic skin-conductance responses (GSR), in addition to their behavioral performance on the comprehension questions. This yielded a rich neurophysiological dataset allowing use to study the impact of background noise on performance, neural processing and physiological responses during realistic VR classroom learning, as well as to test for potential differences between the detrimental effects of intermittent and continuous construction noise.

**Figure 1.**
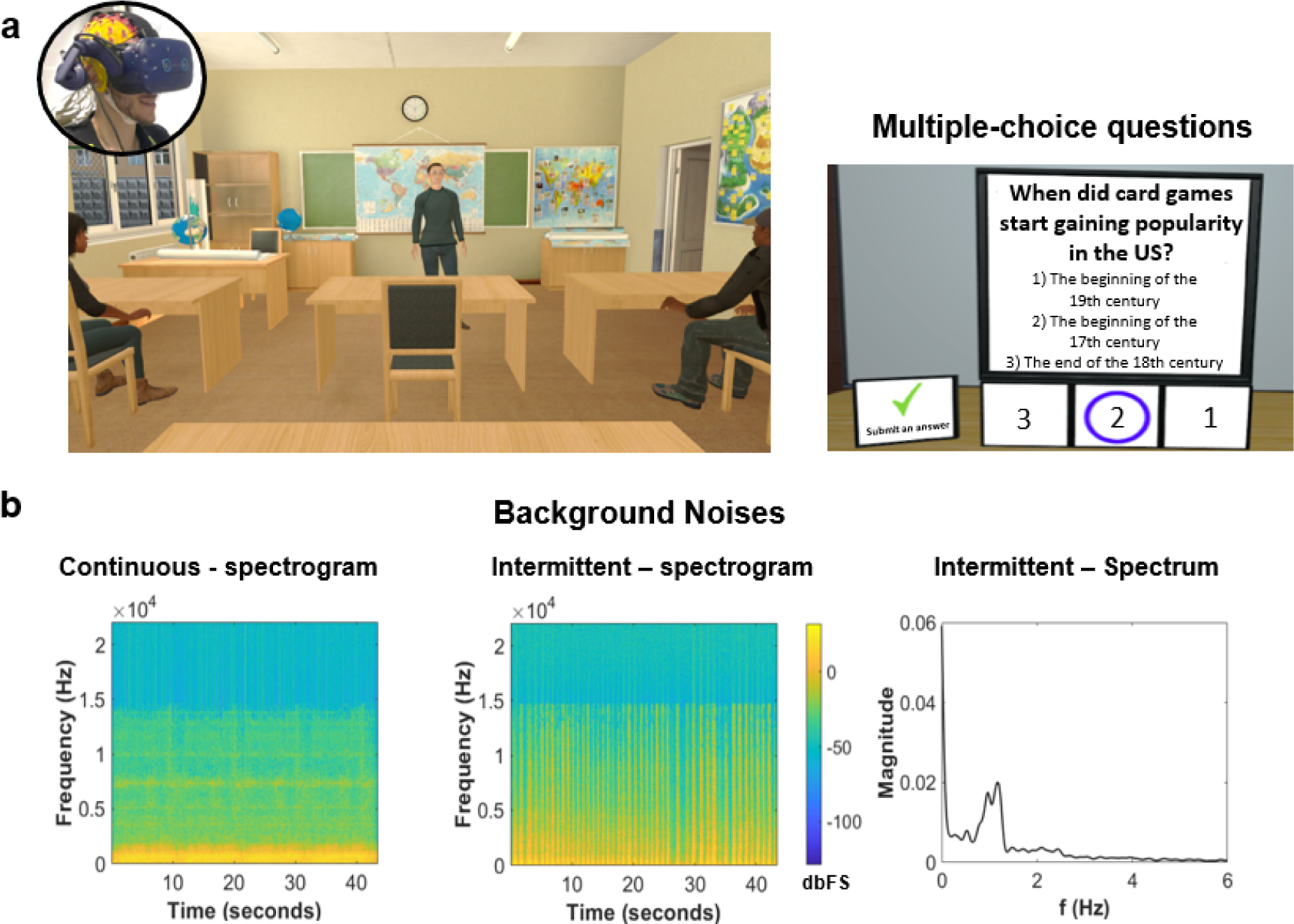
a) Left: Illustration of the Virtual Classroom scenario. The circular inset shows a participant wearing a virtual-reality headset over an EEG cap. Right: example of one multiple-choice question about the teacher content. b) Features of the background noises added to the scene. The left panel depicts the spectrogram of the continuous noise, the middle panel displays the spectrogram of the intermittent noise, and the right panel shows the modulation spectrum of the intermittent noise.

## Methods

### Participants

The experiment included 32 participants, aged between 20 and 39 (mean 24.6 ± 3.8 std; 19 females, 13 males, 25 left-handed). Two participants were excluded from the EEG and GSR analysis due to technical issues and artifacts, and one participant was excluded from all analyses due to a technical issue with randomization in the study design. All participants were native Hebrew speakers with normal hearing and normal or corrected-to-normal vision. None of the subjects had a history of neurological disorders or ADHD (based on self-report). Participants were compensated with either payment or course credit for their participation. The study was approved by the Institutional Review Board committee at Bar-Ilan University, and all participants provided their written consent for participation prior to the experiment.

### The VR Classroom: Design and Stimuli

The VR classroom scene used in this experiment is shown in Figure 1a. The virtual environment was designed using the game engine Unity (https://unity.com/products/unity-engine). It was presented to participants through a Virtual-Reality headset with an embedded eye-tracker (HTC Vive Pro EYE) and audio was presented through in-ear headphones (ER1-14 with disposable foam eartips).

Participants experienced sitting at a desk in the second row of the VR classroom, facing an avatar of a male teacher, standing in front of a blackboard and a wall with maps. Ten additional student avatars sat at the other desks in the classroom. They didn’t talk during the experiment, but their bodies were animated slightly to mimic natural sitting motions.

During the experiment, the speech and body motions of the teacher avatar were controlled, to have him deliver a series of 30 content vignettes (mean duration 42.7 sec ±□5.1). The audio of these content vignettes were segments from an informational podcast, all recorded by the same male speaker, and focused on scientific or historical topics. The audio of these content vignettes was presented using 3D spatial audio (implemented in Unity) as emitting from the teacher’s lips. The teacher’s lip movements were animated to follow the temporal envelope of the speech, and his body motions were animated to capture appropriate emphasis and prosodic cues.

In some trials, background noise was added to the scene, emitting from a virtual location outside the middle-left window of the classroom, which was located approximately 17 meters away from the participant (in virtual space; 3D sound algorithm, Unity). Two different types of noise were used: Continuous noise of air hammers and Intermittent noise of drilling. The intermittent drilling was fairly rhythmic, with bursts of noise every 0.2-1 sec, lasting approximately 660 ms (Figure 1b). These sounds were recorded from a real-life construction site using an iPhone, and were edited to have the same duration (45 sec) and their loudness-level was equated using the software Audacity.

In the VR classroom, the teacher’s speech was presented at a loudness level, that was determined during a training trial. The precise intensity level cannot be measured reliably, since it is affected by the specific positioning of the foam earphone inserts in the participants’ ear, but was roughly between 60-70 dB SPL. The noise stimuli (when present), was presented at the same loudness level as the teacher’s speech, however since it was emitted from a farther spatial location its perceived loudness was attenuated by a factor of 1/4 (-12dB spl), relative to the teacher (Unity Audio Spatializer SDK).

### Experimental procedure

The experiment was conducted in a sound attenuated room. After comfortably fitting the VR headset on the participant, the eye-tracker was calibrated and validated using a standard 9-point calibration procedure (Vive ProEye). Participants were then familiarized with the VR classroom scene, and performed two training trials – one without noise and one with intermitted noise – for training and calibration purposes.

The experiment consisted of 30 trials (∼40 seconds each). In each trial, a different mini-lecture was delivered and participants were instructed to pay attention to the content of the lecture. They were not given any specific instructions regarding other sounds they might hear or about other stimuli in the classroom environment. At the end of each trial, participants were asked four multiple-choice questions regarding the content of the mini-lecture they had just listened to (Figure 1a).

Each mini-lecture was assigned to one of three noise-conditions: Quiet (10 trials), Continuous Noise (10 trials), and Intermitted Noise (10 trials). The allocation of lecture to condition was fully counterbalanced across participants, such that each lecture appeared in each noise-condition in 1/3 of the participants. In addition, the presentation order of the content vignettes was fully randomized across participants.

### EEG, GSR and Eye-Gaze recordings

Neural activity was recorded using 64 active ag-agCl EEG electrodes (BioSemi) and was sampled at 1024Hz. Electrooculographic (EOG) signals were also included in this recording, measured by 3 additional electrodes located above the right eye and on the external side of both eyes.

The Galvanic Skin Response (GSR; detailed description in: Akash et al., 2018) was recorded using two passive Ni hon Kohden electrodes placed on the fingertips of the index and middle fingers of participants’ non-dominant hand. These signals were also sampled by the BioSemi system, and were synchronized to EEG data. Eye-gaze data was recorded using the eye tracker embedded in the VR headset (120 Hz binocular sampling rate), which provides information about the x-y-z location of the gaze, as well as automatic blink-detection, at each time-point. In addition, we defined several a-priori regions of interest (ROIs) as invisible colliders in the environment, using Unity. These included objects such as: ‘Teacher’, ‘Left Board’, ‘Right Board’, ‘Ceiling’, ‘Floor’, ‘Middle Window’, ‘Left Student’, ‘Right Student’ etc. Using a combined calculation of the participants’ head position and direction of the gaze vector, this allowed for automatic classification of the gaze data, describing which ROI the participant was looking at, at each point in time, which was provided as output.

### Behavioral data analysis

To assess participants’ comprehension and memory retention, we analyzed the percentage of correct responses on multiple-choice questions that followed each trial. Responses were compared across conditions using Wilcoxon Signed-Rank to test for differences between the Quiet vs. Noise conditions (averaged across two noise conditions) and for differences between Intermittent vs. Continuous.

### EEG data analysis

EEG preprocessing was performed in MATLAB (MathWorks) using the FieldTrip toolbox (https://www.fieldtriptoolbox.org/). Data were first referenced to the average signal of the mastoids, followed by a band-pass filter at 0.5-40 Hz, de-trended and de-meaned. Then, the data was inspected visually and gross artifacts (that were not eye-movements) were marked and removed. Independent Component Analysis (ICA) was performed to identify and remove components associated with horizontal or vertical eye movements as well as heartbeats. Noisy electrodes that exhibited consistent high-frequency activity (>40Hz) or excessive low-frequency DC drifts, likely due to bad or loose connectivity, were replaced with the weighted average of their neighbours using an interpolation procedure. This was carried out either on the entire dataset or on a per-trial basis, as needed.

#### Speech tracking response (TRF)

The clean data was segmented into trials from the start of the audio until the end of the mini-lecture. An average of 2.03 trials (2.23 std) were removed from each participant due to excessive artifacts. The segmented data were bandpass filtered between 0.8 and 20 Hz using a band-pass filter, and then downsampled to 100 Hz, for computational efficiency. To estimate the neural response to the speech we performed speech-tracking analysis on the data, and estimated Temporal Response Functions (TRF’s) using the mTRF MATLAB toolbox (Crosse et al., 2016). The TRF is a mathematical model that represents a linear transfer function, elucidating the relationship between features of a stimulus (S) and neural response (R) recorded when it is presented. Specifically, we used the broadband envelopes of the speech as stimulus features, which were extracted using an equally spaced filterbank with frequencies ranging from 100 to 10,000 Hz, based on the cochlear frequency map developed by Liberman (1982). The narrowband filtered signals were summed across the frequency bands after rectifying them with the Hilbert transform, resulting in a broadband envelope signal. This envelope was further downsampled to 100 Hz to match the EEG data.

Using the encoding approach, Temporal Response Functions (TRFs) were estimated at different time delays for each EEG channel. These TRFs represent the transfer-function responses to the speaker and were optimized to obtain the most accurate linear combination for predicting the neural response. They can be conceptualized as a linear composition of partially overlapping neural responses to a continuous stimulus, and are therefore analogous to Event Related Potentials (ERPs). As such, the peaks of the TRF can be interpreted in a comparable manner as more traditional ERPs. Thus, encoding models provide a comprehensive and detailed characterization of the neural response to stimuli, revealing both the temporal dynamics and topography of the response across sensors.

TRFs were derived separately for trials in each noise-condition (Quiet, Intermitted, Continuous), including a baseline period and computed over a time range spanning from -150ms (pre-stimulus) to 450ms. A leave-one-out approach was employed to optimize the model, where 29 trials were selected in each iteration for training, and the remaining trial was used for testing. The goodness of fit (predictive power) of the model was determined by calculating the Pearson correlation between the predicted and actual neural response at each sensor. This process was repeated 29 times, with different train-test partitions for each iteration, and the predictive power was averaged over all iterations. To avoid overfitting of the model, a ridge parameter (λ) was selected through a cross-validation process. Since this parameter has a significant impact on the shape and amplitude of the TRF, a common λ value was chosen for all participants to allow for group-level analyses. For each participant, a range of λ values from 10^-2 to 10^4 was tested, and the value that resulted in the highest average predictive power across all channels and participants was selected as the common optimal λ, which was determined to be 100. All results reported in this study utilized this common ridge parameter value.

Before examining the effects of attention, we first assessed which channels exhibited a significantly higher predictive power than chance, using a permutation test. This involved shuffling the pairing between acoustic envelopes (S) and neural data responses (R) across trials, such that the speech-envelope presented in one trial was paired with the neural response recorded in a different trial. This process was repeated 100 times, and an encoding model was optimized for each permutation. We obtained a “max-chance predictive power” null-distribution by selecting the maximum r-value from the grand average across participants for each permutation. EEG channels with r-values exceeding the top 5% of the null distribution were deemed to have a significant speech tracking response.

Next, we tested for differences in the speech tracking response (TRF) and its predictive power across the 3 noise conditions. To assess whether speech tracking was affected by the presence of any type of noise, we compared responses in the Quiet condition vs. the average across the two noise conditions. To test whether the TRFs were affected by the specific type of noise, we further compared TRFs in the Intermitted vs. Continuous condition. In both analyses, we focused on the two observable peaks in the TRFs – a negative peak between 80-110ms and a positive peak between 140-180ms. We performed paired t-tests at each electrode for TRF comparisons and corrected for multiple comparisons using spatio-temporal clustering.

#### Alpha power

To extract the induced alpha-power across trials, we calculated the spectral power density (PSD) using the multitaper fast-fourier transform (FFT) method in Fieldtrip (method ‘mtmfft’, Hanning tapers). The spectral power estimates were averaged across trials in each condition, separately for each channel. We then used the FOOOF algorithm (Donoghue et al., 2020) to decompose the power spectrum density (PSD) into periodic (oscillatory) and aperiodic components. For each individual, we identified the alpha-power peak in the periodic component, as the frequency with the largest averaged amplitude between 6-15Hz in a cluster of pre-defined parietal-occipital electrodes (this band is slightly broader than the canonical alpha-band, but seemed to better capture individual differences in peak-frequencies in the current dataset).

Alpha-power was compared across conditions using Wilcoxon Signed-Rank to test for differences between the Quiet vs. Noise conditions (averaged across two noise conditions) and for differences between Intermittent vs. Continuous.

### Eye tracking data analysis

Data from the eye-tracker, was used to assess the spontaneous gaze-patterns of participants, as they listened to the teacher in the VR classroom, and to test whether participants performed more gaze-shifts away from the teacher in conditions that were noisy. Gaze-data was analyzed in Python using in-house scripts, following recommended procedures for analyzing gaze data in VR (Anderson et al., 2023). We first cleaned gaze-data by removing measurements around blinks (from 100ms before to 200 ms after), and any other data-points that the eye-tracker marked as “missing data”.

To assess gaze-shift patterns, we calculated the Euclidian distance between adjacent data points (using the x,y,z coordinates). ‘Fixations’ were defined as clusters of adjacent time-points, within a radio of less than 0.01 distance from each other, lasting at least 80 ms. ‘Gaze-shifts’ were defined as transitions between Fixation as defined above, i.e., eye-gaze shifts exceeding a distance of 0.01 which were preceded and followed by a stable period of at least 80 ms. Then, each fixation was associated with a particular ROI in the VR classroom, using the center of mass (average x,y,z value) across all data points the fixation, and the mapping between this location and the pre-defined ROIs (colliders) in the VR classroom. This data was used to quantify the average percent of each trial that participants looked at the teacher (target) vs. all other ROIs, and the number of gaze-shifts away from the teacher per trial.

Each of these metrics was compared using Wilcoxon Signed-Rank to test for differences between the Quiet vs. Noise conditions (averaged across two noise conditions) and for differences between Intermittent vs. Continuous.

### GSR data analysis

GSR data analysis was performed using the Matlab-based Ledalab toolbox (Benedek & Kaernbach, 2010). Continuous decomposition analysis (CDA) was used to extract the sequence of tonic and phasic skin-conductance responses (SCR’s) over the duration of each trial for each subject, per condition. For each trial, we computed the average of the phasic driver (SCR), the number of significant phasic SCRs exceeding an amplitude threshold of 0.01 microsiemens (nSCR) and the mean tonic activity. Each of these metrics was compared using Wilcoxon Signed-Rank to test for differences between the Quiet vs. Noise conditions (averaged across two noise conditions) and for differences between Intermittent vs. Continuous. False Discovery Rate (FDR) correction was applied to correct for multiple comparisons.

## Results

### Behavioral results

Participants performed well on the comprehension task with an average accuracy of 91.56 ± 3.509% (std), indicating that participants paid attention to the teacher’s speech as instructed.

A Wilcoxon Signed-Rank test revealed that performance was slightly better in the Quiet condition relative to the two Noise conditions combined [W(30)=148.5, p=0.083; Figure 2]. When directly comparing performance in the Intermittent and Continuous noise conditions, we found a significant difference between them [W(30)=89.5, p=0.047], indicating that participants performed better in the Continuous noise condition compared to the Intermittent noise condition Figure 2.

**Figure 2.**
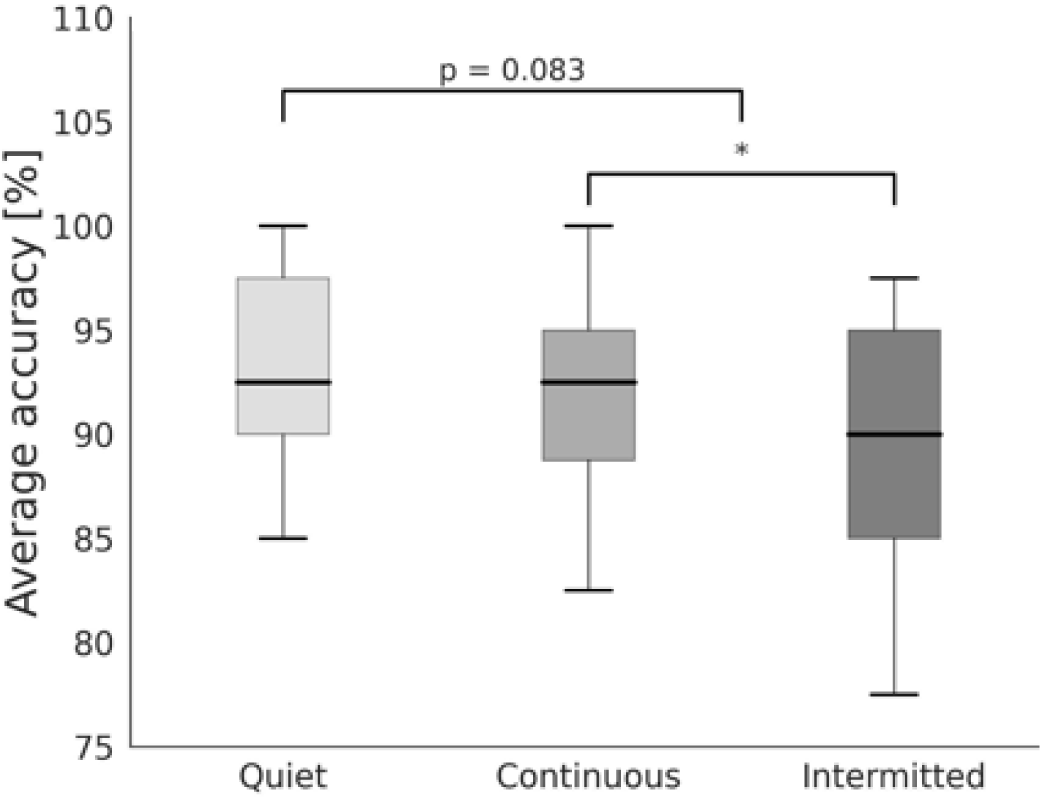
Box plots representing the median behavioral performance on the multiple-choice questions (n = 31) across different conditions. The error bars indicate the standard error of the mean (SEM), and an asterisk (*) indicates a significant difference.

### Post-experiment debriefing

Following the experiment, participants were interviewed about their experience with the different noise conditions. In response to the question *“Did you notice which direction the noise came from?”*, all participants correctly identified that the noise came from the left, indicating that they had all noticed the presence of the noise. When asked *“Did you notice any difference between the types of noises?”*, 64.52% of participants reported that they had noticed that there were two types of noises, one continuous and one intermittent. When asked *“Did the noises bother you or make it difficult to concentrate?”*, 29% of participants reported that the noises did not bother them and even helped them concentrate, 26% reported that the noises sometimes bothered them, but they sometimes got used to them or tried to overcome them, and 45% reported that the noises bothered them and that they preferred the silence condition.

### EEG results

#### Speech tracking response

We performed speech-tracking analysis of the neural responses to speech separately for each condition (Quiet, Intermittent, Continuous). Figure 3a shows the topographical distribution of the predictive power values of the resulting model. Comparison to a null distribution of permuted data revealed clusters of electrodes with significant predictive power in all conditions, primarily over frontal and temporal regions, indicating that a robust speech tracking response could be extracted (p<0.05 corrected; marked in Figure 3a). However, the number of electrodes with a significant speech tracking response was lowest in the Intermittent-Noise condition and was highest in the Quiet condition. In total, the number of electrodes for the Quiet, Continuous, and Intermittent conditions was 24, 15, and 7, respectively.

**Figure 3.**
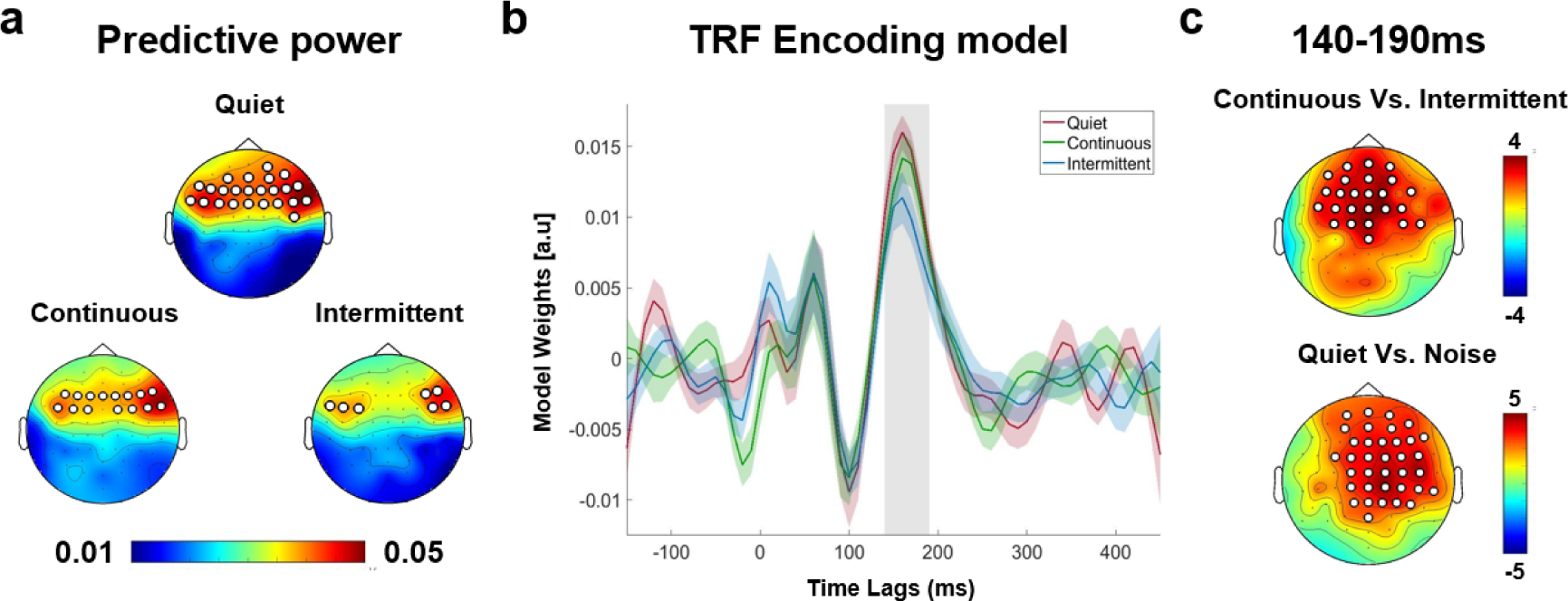
The speech tracking neural response (n=29). a) Topographical distribution of predictive power values for each condition in the speech tracking neural response analysis (n=29). Significant electrode clusters (p<0.05 corrected) compared to the null distribution are marked with white circles b) Temporal Response Functions (TRFs) for each condition. The light gray area represents a significant difference between conditions (p<0.05 corrected) c) Topographical maps showing clusters of significant electrodes (p<0.05 corrected) during the time window of 140-190ms, highlighting differences in TRF amplitudes between the conditions.

Visual inspection of the estimated TRFs showed two prominent peaks – a negative peak around 100ms followed by a positive peak around 150ms (Figure 3b), in line with previous studies. Statistical analysis focused on the TRF amplitudes in time windows surrounding these two peaks (between 80-110 ms and between 140-190 ms). For the early negative peak, we did not find any significant differences between the conditions. However, for the later positive peak, the TRF amplitude was higher in the Quiet condition vs. the two Noise conditions, and was also higher in the Continuous noise condition vs. the Intermittent noise condition (p<0.05, two-tailed cluster correction). The clusters of significant electrodes are marked on the topographical maps in Figure 3c.

We conducted a control analysis to test whether the reduced predictive power and reduced TRF amplitude in the Intermittent condition could be attributed trivially to the additional neural response to the noise itself. Using a multivariate TRF model we tested whether including both the speech-stimulus and the noise-stimulus in the encoding model would improve its predictive power (Crosse et al., 2016). However, we did not find any significant difference between the predictive power of the multivariate model vs. the univariate model (that contained only the speech-stimulus; p=0.33, average across all electrodes), indicating that adding the temporal-modulation of the Intermittent noise did not explain any additional variance in the neural signal.

#### Alpha power

The averaged alpha-power peak between 6-15Hz, calculated across participants and focused on a predefined cluster of parietal-occipital electrodes, is presented in Figure 4a,c. For each participant, we determined the alpha-power peak as the frequency with the highest average amplitude within this frequency range and across trials in different noise conditions.

**Figure 4.**
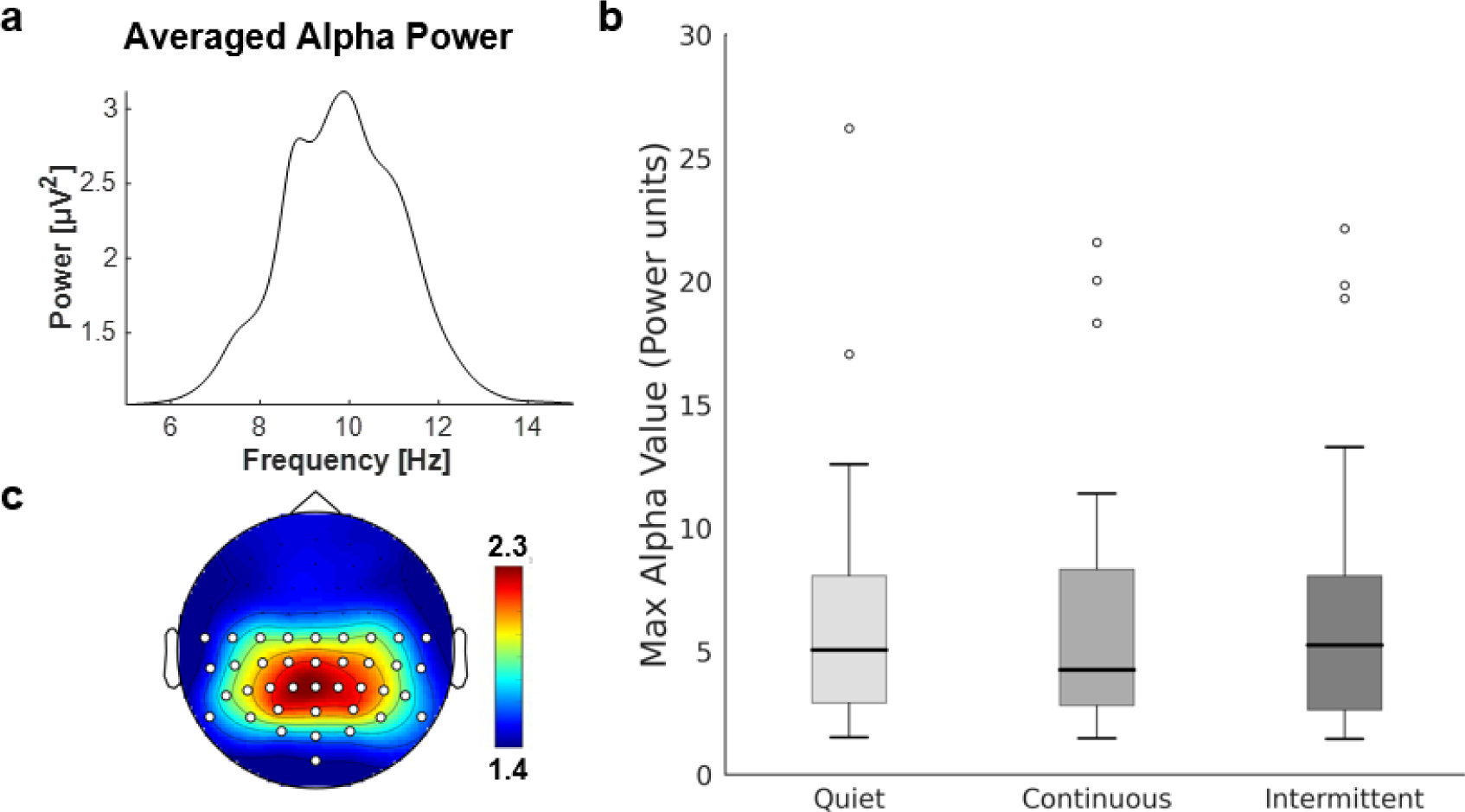
Alpha power results (n=29). a) Averaged alpha-power peak between 6-15Hz focused on a predefined cluster of parietal-occipital electrodes. b) Comparison of the maximum alpha power value across conditions. Outliers represented by circles. c) Topographical distribution of the averaged alpha-power peak, with the cluster of parietal-occipital electrodes marked with white circles.

Wilcoxon Signed-Rank test revealed no differences between the Quiet and Noise conditions [W(28)=183, p=0.46], and a non-significant trend of the difference between the Intermittent and Continuous noise conditions [W(28)=145, p=0.12](Figure 4b). Note, however, that the magnitude of alpha power varied substantially across participants, and did not follow a normal distribution.

### Eye tracking results

We analyzed the proportion of each trial that participants spent looking at the teacher and at different areas in the VR classroom, as well as the number of gaze-shifts away from the teacher. As shown in Figure 5a, participants spent most of the time looking at the teacher [average 66.86 ± 21.49 % of each trial]. When not looking at the teacher, the next most popular places to look at were the boards behind the teacher, to the right and to the left. Glances were occasionally directed to the floor, the windows and other student-avatars.

**Figure 5.**
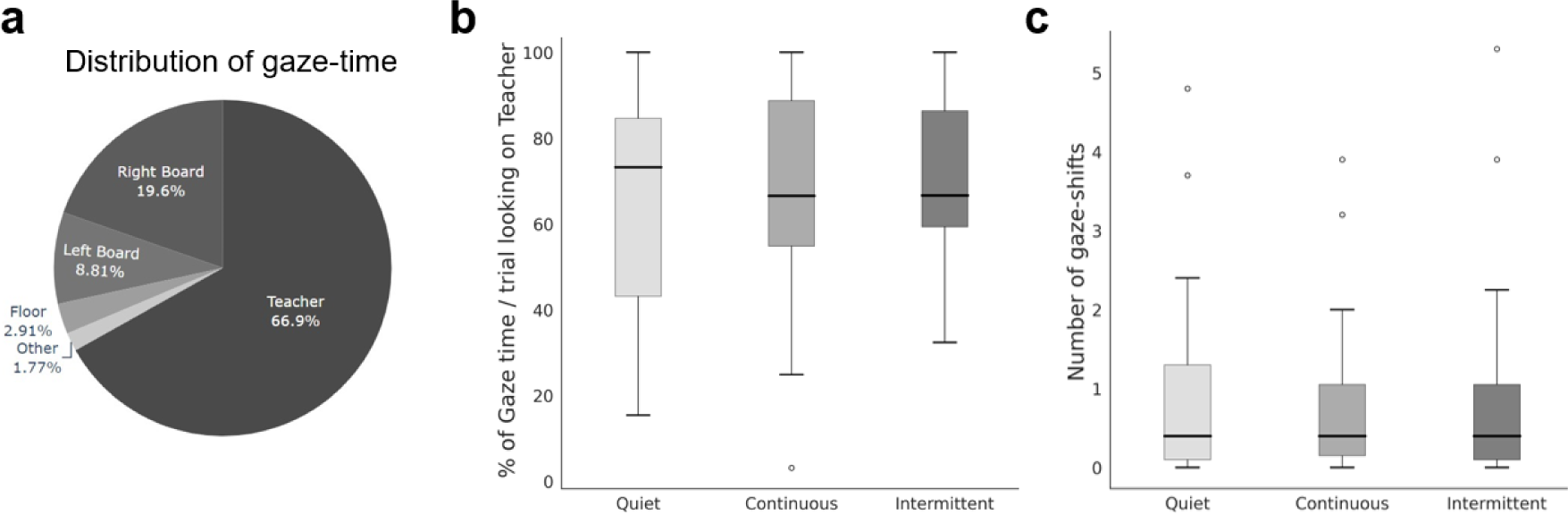
Eye-tracking results (n=31). a) Pie chart represents the distribution of eye gaze to areas of the classroom (using predefined ROIs). b) Box plots represent the percent of time spent looking at the teacher in different noise conditions c) Box plots represent the number of gaze shifts away from the teacher between conditions. In all figures the circles represent outliers.

We did not find significant differences between conditions in any of the eye-gaze metrics, including: the percent of time spent looking at the teacher in the different noise conditions [Quiet vs. Noise: W(30) = 202, p = 0.53; Intermittent vs. Continuous: W(30) = 151, p = 0.15; Figure 5b] or in the number of gaze-shifts performed away from the teacher [Quiet vs. Noise: W(30) = 200.5, p = 0.95; Intermittent vs. Continuous: W(30) = 172, p = 0.92; Figure 5c].

### GSR results

For each participant, we collected various metrics of Galvanic Skin Response (GSR), including the level of Tonic response, the number of Phasic responses (nSCR) and the mean amplitude of Phasic responses (SCR). We conducted Wilcoxon Signed-Rank tests to compare the Quiet condition with the two Noise conditions and the Intermittent condition with the Continuous condition for each metric. After applying the False Discovery Rate (FDR) correction, we found no significant differences between the Quiet and Noise conditions in any of the metrics (all ps > 0.5). Furthermore, there was no significant difference observed in the SCR between the Intermittent and Continuous conditions [SCR: W(28) = 161, p = 0.23]. However, we did find a significant difference in their nSCR and Tonic responses, with a higher number of Phasic responses and a non-significant trend towards higher Tonic response in the Intermittent condition [W(28) = 97.5, p = 0.048 and W(28) = 132.5, p = 0.103, respectively; refer to Figure 6].

**Figure 6.**
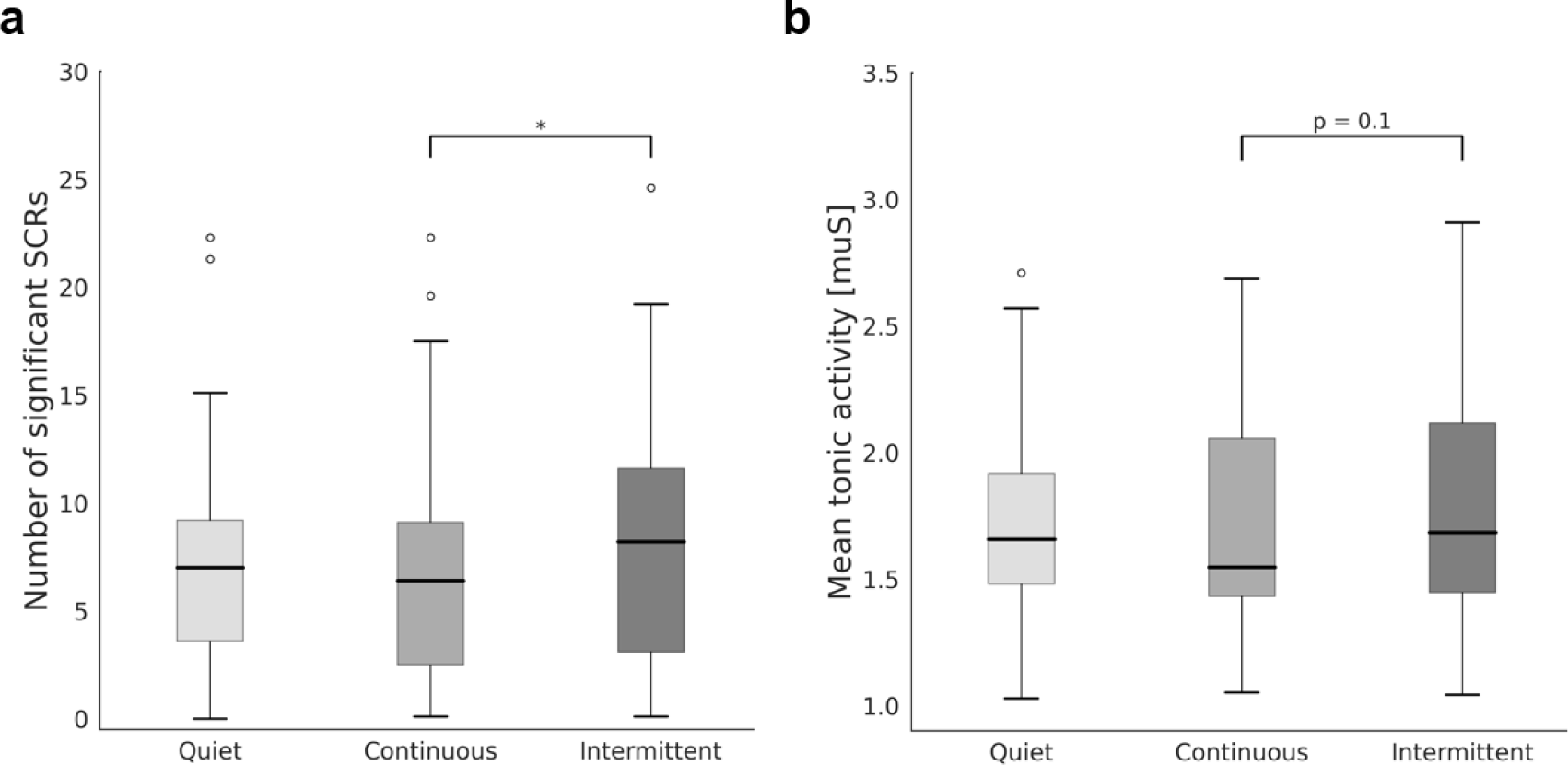
Galvanic Skin Response (GSR) results (n=29). a) Box plots represent the number of significant SCRs exceeding an amplitude threshold of 0.01 microsiemens in different noise conditions. An asterisk (*) indicates a significant difference (p<0.05). circles represent outliers. b) Box plots represent the mean tonic activity in microsiemens between conditions. circles represent outliers.

## Discussion

In this study, we collected a rich neurophysiological dataset to investigate the impact of background construction noise during realistic learning in a VR classroom. As expected, the presence of noise had an overall detrimental effect, leading to reduced behavioral performance and reduced neural tracking of the teacher’s speech, as well as altered physiological responses. These disruptive neurophysiological effects were more pronounced for intermittent vs. continuous noise, indicating that the temporal structure of noise contributes to its negative impact, above and beyond loudness level.

### The disruptive influence of noise

The difficulty to understand speech under noisy conditions is attributed to three factors: masking, selective attention and listening effort. *Masking* is the acoustic interference and spectral overlap of background noise on target speech. This poses a perceptual challenge of segregating the overlapping auditory inputs and extracting a clear representation of the target speech (Darwin, 2008; Heinrich et al., 2008; Mattys et al., 2012; Peelle, 2018). In attempt to overcome the perceptual challenge of masking, the cognitive system employs top-down *selective attention* to actively amplify the target speech above and beyond the background noise (Agmon et al., 2022; Lee et al., 2014; Shinn-Cunningham & Best, 2008). The combined effects of masking and selective attention result in heightened cognitive demand requiring increased *listening effort* in order to process speech in noise (Eckert et al., 2016; Peelle, 2018). This heightened cognitive load redirects attentional resources from other tasks, adversely affecting encoding processes and diminishing success in remembering and learning new material (Eckert et al., 2016; Heinrich et al., 2008). Prolonged cognitive exertion can also cause feelings of effort, exhaustion, and fatigue (Eckert et al., 2016).

The neurophysiological effects of noise observed here during VR learning, likely reflect the consequences of these three processes and the overall increased cognitive effort for processing speech in noise. The most direct consequence of a noisy environment is reduced behavioral performance, a finding that is found repeatedly in lab-based and field work and in the current study (Darwin, 2008; Dubbelboer & Houtgast, 2007; Sala & Rantala, 2016). We show that this behavioral cost is accompanied by two neurophysiological costs: reduced neural tracking of the target speech, found in both noise conditions, and increased skin-conductance, observed here in the presence of intermittent noise. We now turn to discuss these neurophysiological results and their implications for understanding the effects of noise on speech processing under realistic conditions.

### Reduced neural tracking of speech in noise

Neural speech tracking reflects the internal representation of incoming speech in auditory and language-related brain regions. Although this response is driven by speech acoustics, it is known to be modulated by top-down processes including selective attention (Agmon et al., 2022; Har-shai Yahav & Zion Golumbic, 2021; Kaufman & Zion Golumbic, 2023; O’Sullivan et al., 2015; Rimmele et al., 2015; Zion Golumbic et al., 2013) and listening effort (Dimitrijevic et al., 2019; Song & Iverson, 2018), as well as linguistic factors such as intelligibility (Broderick et al., 2018; Di Liberto et al., 2018; Lesenfants et al., 2019; Verschueren et al., 2020). The reduced neural tracking of the teacher’s speech found here in the presence of noise is consistent with previous studies reporting a decrease in speech tracking as noise levels increase (Kong et al., 2015; Vanthornhout et al., 2018; Zou et al., 2019), particularly at intermediate and loud intensities (Ding & Simon, 2013; Hauswald et al., 2022; Yasmin et al., 2023). These effects likely reflect the combined consequences of acoustic masking, reduced intelligibility, reduced attention levels and increased listening effort.

In analyzing the time-course of the speech-tracking response (TRF), we found that noise primarily impacted the mid-latency positive peak response around ∼150ms, but not the earlier negative peak at 100ms which is associated with low-level acoustic encoding (Salmelin, 2007). This is consistent with several recent studies that report correlations between mid-latency TRF components and intelligibility, particularly under noisy conditions (Broderick et al., 2018; Chen et al., 2023; Verschueren et al., 2020). Other studies reported modulation of mid-latency TRF components for degraded vs. clear speech (Kraus et al., 2021; Strauß et al., 2013), for background vs. target speech (Kaufman & Zion Golumbic, 2023; Orf et al., 2023) and for speech in an unfamiliar vs. familiar voice (Har-shai Yahav et al., 2023). Hence, the latency of these effects suggests that in the current dataset, noise did not necessarily impact the earliest levels of acoustic encoding of the speech, but rather affected more downstream processes related to its intelligibility or semantic content.

Reduced speech tracking was observed in both noise conditions, but was worse in the presence of intermittent relative to continuous noise. Our control analysis (use of a multivariate TRF model) confirmed that this was not simply due to the additional neural response evoked by the noise itself, but reflects true reduction in speech tracking response. This pattern converges with the behavioral results, which were also worse in the intermittent noise condition, indicating that intermittent is more disruptive to speech processing than continuous noise, despite their equal intensity (see discussion below).

### Heightened Skin Conductance with Intermittent Noise

Not only was performance worse and speech tracking reduced in the intermittent noise condition, but in this condition we also found an increase in skin conductance, characterized by more frequent phasic responses and a trend toward higher tonic responses, relative to the quiet and continuous noise conditions. These physiological responses reflect heightened activation of the sympathetic nervous system, which is known to respond to salient stimuli, both task-relevant and in the background (Brown et al., 2023; Dawson et al., 1989; Filion et al., 1991; Mueller-Pfeiffer et al., 2014), and is also engaged in conditions requiring high listening effort (Borghini & Hazan, 2018; Mackersie & Cones, 2011; et al., 2017).

Accordingly, the higher skin-conductance responses observed here in the intermittent noise condition suggest that this condition required higher listening effort and/or that the frequent onsets and offsets in this condition drive the system into a heightened state of excitability. Conversely, continuous noise did not lead to a change in skin-conductance metrics, which is in line with previous studies showing stable skin-conductance levels across varying levels of continuous noise (Mackersie et al., 2015) and for ambient noise (Alvarsson et al., 2010); but see caveat by (Cvijanović et al., 2017). This result converges with the behavioral and neural findings to suggest that intermittent background noise is more disruptive and bothersome to listeners than continuous noise.

### Why is intermittent noise more disruptive?

When setting up this experiment, we were agnostic as to which type of noise would be more disruptive, since theoretical predictions could be made in either direction. The glimpsing model, which suggests that processing speech in noise relies on ‘listening in the gaps’ (Cooke, 2006), predicts that intermittent noise would be easier to deal with, relative to continuous noise at the same intensity, due to the presence of gaps. Conversely, theories of neural adaptation predict that continuous noise would undergo adaptation/habituation and therefore would be less disruptive than intermittent noise (Thompson & Spencer, 1966). The current data support the latter perspective, suggesting that the stationary property of continuous noise promotes the adaptation/habituation of its encoding in auditory cortex, leading to reduced masking and ultimately to reduced listening effort. Similar habituation effects of background noise have been demonstrated in previous studies. For example, Bell et al. (2012) found that the interference of background noise decreases significantly after only 45 seconds of exposure. Other studies have also shown noise adaptation in speech recognition tasks (Marrufo-Pérez et al., 2022).

Conversely, the frequent onsets and offsets of intermittent noise make it less adaptable and therefore more disruptive. One way to think about intermittent noise is as a sequence of transient events, each of which can elicit an evoked response and/or a phase-reset in the auditory system, preventing neural adaptation to the background noise (Doelling et al., 2019; Haig & Gordon, 1998; Henry & Obleser, 2012; Hodapp & Grimm, 2021; Ten Oever et al., 2017; Uther et al., 2003). Interestingly, although the intermittent noise was relatively rhythmic, its temporal predictability did not necessarily assist in its suppression, at least relative to continuous noise (Makov et al., 2017).

Regarding the glimpsing model of speech in noise perception, the current results suggest that ‘listening in the gaps’ did not facilitate speech processing in our VR classroom. However, it is important to note that the empirical findings that gave rise to the glimpsing model were obtained under different conditions and stimuli than those tested here. In most studies, the speech stimuli consist of single syllables/words or simple sentences, tasks consist of word-by-word recognition, and the noises used are artificial and highly controlled in their spectro-temporal properties (Buss et al., 2003, 2004; Cooke, 2006; Fogerty et al., 2018; Howard-Jones & Rosen, 1993; Kidd et al., 2016; Miller & Licklider, 1950). In contrast, in the current setup we used long segments of continuous speech presented in a realistic audiovisual context, we tested its overall comprehension rather than word-for-word recognition, and used naturally-recorded construction noise that is less controlled and perhaps more abrasive its acoustic properties than the modulated noises used in glimpsing studies (Fogerty et al., 2018). These design choices are part of our attempt to study speech processing in noise under more ecologically-relevant contexts (Brown et al., 2023; Deringer & Hanley, 2021; Parsons, 2015; Risko et al., 2016; Shavit-Cohen & Zion Golumbic, 2019). However, they also probably account for the discrepancies in findings relative to more traditional psychoacoustic experiments, and raise the possibility that while the glimpsing model may hold true for local perception of speech-elements, intermittent noise remains highly disruptive and incurs high cognitive costs when attempting to focus ones’ attention on continuous speech for long periods of time. That said, since in the current study we only tested two specific noise stimuli, these theoretical notions should be further tested in follow-up studies using a wider variety of noises differing in loudness, modulation frequency and depth, temporal structure, and semantic properties.

### Metrics not affected by noise: alpha power and gaze-shifts

Two of the neurophysiological metrics assessed here did not show any modulation by the presence of noise, despite theoretical predictions that they might: neural alpha oscillation and spontaneous gaze-shifts.

Alpha oscillations have been proposed to play an important role in selective attention and cognitive effort, and alpha power is generally found to be reduced under conditions that require high attention and/or cognitive effort (Antonenko et al., 2010; Ehrhardt et al., 2022; Foxe & Snyder, 2011). However, in the context of processing speech in noise, mixed results have been reported. While some have found enhanced alpha power when hearing degraded speech (Becker et al., 2013; Obleser et al., 2012; Wöstmann et al., 2015), others have found the opposite effect (Hauswald et al., 2022; Miles et al., 2017). These discrepancies in empirical findings resonate with the opposing theoretical hypotheses regarding the functional role of alpha oscillations, and whether they plays a role in inhibiting irrelevant information (Foxe & Snyder, 2011; Jensen & Mazaheri, 2010; Strauß et al., 2014) or in facilitating information processing and working memory (Palva & Palva, 2011). Since in the current study we did not find any significant effects of noise on alpha power (although there was a trend toward decreased alpha power the intermittent noise conditions), our data cannot shed light on this debate. However, the conflicting past results and the current null results do highlight the complexity of linking a particular neural metric to specific cognitive or perceptual process (Lozano-Soldevilla, 2018; Schneider et al., 2022).

The second metric that was not affected by noise in the VR classroom were spontaneous gaze-shifts. One of the innovations of using VR is that individuals are free to move their gaze around the virtual environment and are not constrained by a fixation point as in many traditional experiments. The ability to move your eyes freely, is thought to be a central mechanism in controlling and focusing attention (Craighero & Rizzolatti, 2005). Past VR studies have shown that when given this freedom, some participants perform frequent spontaneous gaze-shifts around the environment, while others keep their eyes fixed on the target speaker (Brown et al., 2023; Shavit-Cohen & Zion Golumbic, 2019). When processing audiovisual speech in noise, past studies have shown people tend to focus their gaze more intently on speaker’s face, and particularly on the mouth area, ostensibly to utilize visual cues to overcome the acoustic degradation of the speech (Buchan et al., 2008; Król, 2018; Šabić et al., 2020). Accordingly, we might expect to see reduced gaze-shifts in the noisy vs. quiet conditions, reflecting an attempt to compensate for the acoustic degradation. However, this was not the case. While we replicate the large individual differences in the frequency of spontaneous gaze-shifts reported previously (Brown et al., 2023; Shavit-Cohen & Zion Golumbic, 2019), this was not affected by the presence or type of noise. One possibility is that visual cues were less helpful for speech-processing in the VR classroom due to the participants ‘virtual distance’ from the teacher relative to previous studies where the video of a target speaker is presented full-screen (Aparicio et al., 2017; Šabić et al., 2020). Alternatively, it is possible that the large individual differences in the frequency of gaze-shifts precludes observing group-level effects of noise. We hope to further investigate the dynamics of gaze-shifts in noisy environments using VR and individual differences in future studies.

### Conclusions: Toward more ecological research of speech processing

The present study advances our understanding of the neurophysiological mechanism involved in processing continuous speech under realistically noisy conditions. Use a novel VR classroom platform, we were able to emulate the type of speech and noise stimuli, the task demands and the perceptual complexity of real-life situations, enhancing the ecological validity cognitive neuroscientific research (Deringer & Hanley, 2021; Parsons, 2015; Risko et al., 2016). Our findings demonstrate the multifaceted effects of realistic background noise on listeners, affecting behavior, neural activity, and physiological responses. While not exhaustive, by comparing two types of noises we found that intermittent construction noise was more disruptive to listeners than continuous noise. These findings invite a more dimensional approach to investigating how the spectro-temporal features of realistic noise features affect performance and emphasize the value of studying speech perception in realistic noisy environments using multimodal measurements of neural, ocular, and physiological responses.

## Bibliography

Adams, R., Finn, P., Moes, E., Flannery, K., & Rizzo, A. (2009). Distractibility in Attention/Deficit/Hyperactivity Disorder (ADHD): the virtual reality classroom. Child Neuropsychology□: A Journal on Normal and Abnormal Development in Childhood and Adolescence, 15(2), 120–135. 10.1080/09297040802169077

Agmon, G., Yahav, P. H. S., Ben-Shachar, M., & Golumbic, E. Z. (2022). Attention to speech: mapping distributed and selective attention systems. Cerebral Cortex, 32(17), 3763–3776. 10.1093/CERCOR/BHAB446

Akash, K., Hu, W. L., Jain, N., & Reid, T. (2018). A classification model for sensing human trust in machines using EEG and GSR. ACM Transactions on Interactive Intelligent Systems, 8(4). 10.1145/3132743

Alvarsson, J. J., Wiens, S., & Nilsson, M. E. (2010). Stress Recovery during Exposure to Nature Sound and Environmental Noise. International Journal of Environmental Research and Public Health 2010, Vol. 7, Pages 1036-1046, 7(3), 1036–1046. 10.3390/IJERPH7031036

Anderson, N. C., Bischof, W. F., & Kingstone, A. (2023). Eye Tracking in Virtual Reality. Current Topics in Behavioral Neurosciences. 10.1007/7854_2022_409

Antonenko, P., Paas, F., Grabner, R., & van Gog, T. (2010). Using Electroencephalography to Measure Cognitive Load. Educational Psychology Review, 22(4), 425–438. 10.1007/S10648-010-9130-Y/FIGURES/3

Aparicio, M., Peigneux, P., Charlier, B., Balériaux, D., Kavec, M., & Leybaert, J. (2017). The neural basis of speech perception through lipreading and manual cues: Evidence from deaf native users of cued speech. Frontiers in Psychology, 8(MAR), 230872. 10.3389/FPSYG.2017.00426/BIBTEX

Astolfi, A., Puglisi, G. E., Murgia, S., Minelli, G., Pellerey, F., Prato, A., & Sacco, T. (2019). Influence of Classroom Acoustics on Noise Disturbance and Well-Being for First Graders. Frontiers in Psychology, 10, 482200. 10.3389/FPSYG.2019.02736/BIBTEX

Banbury, S., & Berry, D. C. (1997). Habituation and Dishabituation to Speech and Office Noise. Journal of Experimental Psychology: Applied, 3(3), 181–195. 10.1037/1076-898X.3.3.181

Becker, R., Pefkou, M., Michel, C. M., & Hervais-Adelman, A. G. (2013). Left temporal alpha-band activity reflects single word intelligibility. Frontiers in Systems Neuroscience, 7(DEC), 68736. 10.3389/FNSYS.2013.00121/BIBTEX

Bell, R., Röer, J. P., Dentale, S., & Buchner, A. (2012). Habituation of the irrelevant sound effect: Evidence for an attentional theory of short-term memory disruption. Journal of Experimental Psychology: Learning Memory and Cognition, 38(6), 1542–1557. 10.1037/A0028459

Benedek, M., & Kaernbach, C. (2010). A continuous measure of phasic electrodermal activity. Journal of Neuroscience Methods, 190(1), 80–91. 10.1016/J.JNEUMETH.2010.04.028

Borghini, G., & Hazan, V. (2018). Listening effort during sentence processing is increased for non-native listeners: A pupillometry study. Frontiers in Neuroscience, 12(MAR), 325149. 10.3389/FNINS.2018.00152/BIBTEX

Broderick, M. P., Anderson, A. J., Di Liberto, G. M., Crosse, M. J., & Lalor, E. C. (2018). Electrophysiological Correlates of Semantic Dissimilarity Reflect the Comprehension of Natural, Narrative Speech. Current Biology, 28(5), 803–809.e3. 10.1016/J.CUB.2018.01.080

Brown, A., Pinto, D., Burgart, K., Zvilichovsky, Y., & Zion-Golumbic, E. (2023). Neurophysiological Evidence for Semantic Processing of Irrelevant Speech and Own-Name Detection in a Virtual Café. Journal of Neuroscience, 43(27), 5045–5056. 10.1523/JNEUROSCI.1731-22.2023

Buchan, J. N., Paré, M., & Munhall, K. G. (2008). The effect of varying talker identity and listening conditions on gaze behavior during audiovisual speech perception. Brain Research, 1242, 162–171. 10.1016/J.BRAINRES.2008.06.083

Buss, E., Hall, J. W., Grose, J. H., & Iii, J. W. H. (2004). Spectral integration of synchronous and asynchronous cues to consonant identification. The Journal of the Acoustical Society of America, 115(5), 2278–2285. 10.1121/1.1691035

Buss, E., Wall, J. W., Grose, J. H., & Iii, J. W. W. (2003). Effect of amplitude modulation coherence for masked speech signals filtered into narrow bands. The Journal of the Acoustical Society of America, 113(1), 462–467. 10.1121/1.1528927

Chen, Y. P., Schmidt, F., Keitel, A., Rösch, S., Hauswald, A., & Weisz, N. (2023). Speech intelligibility changes the temporal evolution of neural speech tracking. NeuroImage, 268, 119894. 10.1016/J.NEUROIMAGE.2023.119894

Cooke, M. (2006). A glimpsing model of speech perception in noise. The Journal of the Acoustical Society of America, 119(3), 1562–1573. 10.1121/1.2166600

Craighero, L., & Rizzolatti, G. (2005). The Premotor Theory of Attention. Neurobiology of Attention, 181–186. 10.1016/B978-012375731-9/50035-5

Crosse, M. J., Di Liberto, G. M., Bednar, A., & Lalor, E. C. (2016). The multivariate temporal response function (mTRF) toolbox: A MATLAB toolbox for relating neural signals to continuous stimuli. Frontiers in Human Neuroscience, 10(NOV2016), 219245. 10.3389/FNHUM.2016.00604/BIBTEX

Cvijanović, N., Kechichian, P., Janse, K., & Kohlrausch, A. (2017). Effects of noise on arousal in a speech communication setting. Speech Communication, 88, 127–136. 10.1016/J.SPECOM.2017.02.001

Darwin, C. J. (2008). Listening to speech in the presence of other sounds. In Philosophical Transactions of the Royal Society B: Biological Sciences (Vol. 363, Issue 1493, pp. 1011–1021). Royal Society. 10.1098/rstb.2007.2156

Dawson, M. E., Filion, D. L., & Schell, A. M. (1989). Is Elicitation of the Autonomic Orienting Response Associated With Allocation of Processing Resources? Psychophysiology, 26(5), 560–572. 10.1111/J.1469-8986.1989.TB00710.X

Deringer, S. A., & Hanley, A. (2021). Virtual Reality of Nature Can Be as Effective as Actual Nature in Promoting Ecological Behavior. Ecopsychology, 13(3), 219–226. 10.1089/ECO.2020.0044/ASSET/IMAGES/LARGE/ECO.2020.0044_FIGURE4.JPEG

Di Liberto, G. M., Lalor, E. C., & Millman, R. E. (2018). Causal cortical dynamics of a predictive enhancement of speech intelligibility. NeuroImage, 166, 247–258. 10.1016/J.NEUROIMAGE.2017.10.066

Dimitrijevic, A., Smith, M. L., Kadis, D. S., & Moore, D. R. (2019). Neural indices of listening effort in noisy environments. Scientific Reports 2019 9:1, 9(1), 1–10. 10.1038/s41598-019-47643-1

Ding, N., & Simon, J. Z. (2013). Adaptive Temporal Encoding Leads to a Background-Insensitive Cortical Representation of Speech. Journal of Neuroscience, 33(13), 5728– 5735. 10.1523/JNEUROSCI.5297-12.2013

Doelling, K. B., Florencia Assaneo, M., Bevilacqua, D., Pesaran, B., & Poeppel, D. (2019). An oscillator model better predicts cortical entrainment to music. Proceedings of the National Academy of Sciences of the United States of America, 116(20), 10113–10121. 10.1073/PNAS.1816414116/-/DCSUPPLEMENTAL

Donoghue, T., Haller, M., Peterson, E. J., Varma, P., Sebastian, P., Gao, R., Noto, T., Lara, A. H., Wallis, J. D., Knight, R. T., Shestyuk, A., & Voytek, B. (2020). Parameterizing neural power spectra into periodic and aperiodic components. Nature Neuroscience 2020 23:12, 23(12), 1655–1665. 10.1038/s41593-020-00744-x

Dubbelboer, F., & Houtgast, T. (2007). A detailed study on the effects of noise on speech intelligibility. The Journal of the Acoustical Society of America, 122(5), 2865–2871. 10.1121/1.2783131

Eckert, M. A., Teubner-Rhodes, S., & Vaden, K. I. (2016). Is Listening in Noise Worth It? The Neurobiology of Speech Recognition in Challenging Listening Conditions. Ear and Hearing, 37(Suppl 1), 101S. 10.1097/AUD.0000000000000300

Ehrhardt, N. M., Fietz, J., Kopf-Beck, J., Kappelmann, N., & Brem, A. K. (2022). Separating EEG correlates of stress: Cognitive effort, time pressure, and social-evaluative threat. European Journal of Neuroscience, 55(9–10), 2464–2473. 10.1111/EJN.15211

Eqlimi, E., Bockstael, A., De Coensel, B., Schönwiesner, M., Talsma, D., & Botteldooren, D. (2020). EEG Correlates of Learning From Speech Presented in Environmental Noise. Frontiers in Psychology, 11, 542295. 10.3389/FPSYG.2020.01850/BIBTEX

Filion, D. L., Dawson, M. E., Schell, A. M., & Hazlett, E. A. (1991). The Relationship Between Skin Conductance Orienting the Allocation of Processing Resources. Psychophysiology, 28(4), 410–424. 10.1111/J.1469-8986.1991.TB00725.X

Fogerty, D., Carter, B. L., Healy, E. W., & Jjl, [. (2018). Glimpsing speech in temporally and spectro-temporally modulated noise. The Journal of the Acoustical Society of America, 143(5), 3047–3057. 10.1121/1.5038266

Foxe, J. J., & Snyder, A. C. (2011). The role of alpha-band brain oscillations as a sensory suppression mechanism during selective attention. Frontiers in Psychology, 2(JUL), 10747. 10.3389/FPSYG.2011.00154/BIBTEX

Haig, A. R., & Gordon, E. (1998). EEG alpha phase at stimulus onset significantly affects the amplitude of the P3 ERP component. International Journal of Neuroscience, 93(1–2), 101–115. 10.3109/00207459808986416

Har-shai Yahav, P., Sharaabi, A., & Zion Golumbic, E. (2023). The effect of voice familiarity on attention to speech in a cocktail party scenario. Cerebral Cortex. 10.1093/CERCOR/BHAD475

Har-shai Yahav, P., & Zion Golumbic, E. (2021). Linguistic processing of task-irrelevant speech at a cocktail party. ELife, 10. 10.7554/ELIFE.65096

Hauswald, A., Keitel, A., Chen, Y. P., Rösch, S., & Weisz, N. (2022). Degradation levels of continuous speech affect neural speech tracking and alpha power differently. The European Journal of Neuroscience, 55(11–12), 3288. 10.1111/EJN.14912

Heinrich, A., Schneider, B. A., & Craik, F. I. M. (2008). Investigating the influence of continuous babble on auditory short-term memory performance. Quarterly Journal of Experimental Psychology (2006), 61(5), 735–751. 10.1080/17470210701402372

Henry, M. J., & Obleser, J. (2012). Frequency modulation entrains slow neural oscillations and optimizes human listening behavior. Proceedings of the National Academy of Sciences of the United States of America, 109(49), 20095–20100. 10.1073/PNAS.1213390109/SUPPL_FILE/PNAS.201213390SI.PDF

Hodapp, A., & Grimm, S. (2021). Neural signatures of temporal regularity and recurring patterns in random tonal sound sequences. The European Journal of Neuroscience, 53(8), 2740–2754. 10.1111/EJN.15123

Holleman, G. A., Hooge, I. T. C., Kemner, C., & Hessels, R. S. (2020). The ‘Real-World Approach’ and Its Problems: A Critique of the Term Ecological Validity. Frontiers in Psychology, 11, 529490. 10.3389/FPSYG.2020.00721/BIBTEX

Howard-Jones, P. A., & Rosen, S. (1993). Uncomodulated glimpsing in ‘“checkerboard”’ noise. The Journal of the Acoustical Society of America, 93(5), 2915–2922. 10.1121/1.405811

Hygge, S., Boman, E., & Enmarker, I. (2003). The effects of road traffic noise and meaningful irrelevant speech on different memory systems. Scandinavian Journal of Psychology, 44(1), 13–21. 10.1111/1467-9450.00316

Jensen, O., & Mazaheri, A. (2010). Shaping functional architecture by oscillatory alpha activity: Gating by inhibition. Frontiers in Human Neuroscience, 4, 2008. 10.3389/FNHUM.2010.00186/BIBTEX

Kaufman, M., & Zion Golumbic, E. (2023). Listening to two speakers: Capacity and tradeoffs in neural speech tracking during Selective and Distributed Attention. NeuroImage, 270, 119984. 10.1016/J.NEUROIMAGE.2023.119984

Kidd, G., Mason, C. R., Swaminathan, J., Roverud, E., Clayton, K. K., & Best, V. (2016). Determining the energetic and informational components of speech-on-speech masking. The Journal of the Acoustical Society of America, 140(1), 132–144. 10.1121/1.4954748

Klatte, M., Hellbrück, J., Seidel, J., & Leistner, P. (2010). Effects of Classroom Acoustics on Performance and Well-Being in Elementary School Children: A Field Study. Environment and Behavior, 42(5), 659–692. 10.1177/0013916509336813

Koelewijn, T., Zekveld, A. A., Festen, J. M., & Kramer, S. E. (2012). Pupil dilation uncovers extra listening effort in the presence of a single-talker masker. Ear and Hearing, 33(2), 291–300. 10.1097/AUD.0B013E3182310019

Kong, Y. Y., Somarowthu, A., & Ding, N. (2015). Effects of Spectral Degradation on Attentional Modulation of Cortical Auditory Responses to Continuous Speech. JARO - Journal of the Association for Research in Otolaryngology, 16(6), 783–796. 10.1007/S10162-015-0540-X/FIGURES/7

Kozou, H., Kujala, T., Shtyrov, Y., Toppila, E., Starck, J., Alku, P., & Näätänen, R. (2005). The effect of different noise types on the speech and non-speech elicited mismatch negativity. Hearing Research, 199(1–2), 31–39. 10.1016/J.HEARES.2004.07.010

Kraus, F., Tune, S., Ruhe, A., Obleser, J., & Wöstmann, M. (2021). Unilateral Acoustic Degradation Delays Attentional Separation of Competing Speech. Trends in Hearing, 25. 10.1177/23312165211013242

Król, M. E. (2018). Auditory noise increases the allocation of attention to the mouth, and the eyes pay the price: An eye-tracking study. PloS One, 13(3). 10.1371/JOURNAL.PONE.0194491

Lee, A. K. C., Larson, E., Maddox, R. K., & Shinn-Cunningham, B. G. (2014). Using neuroimaging to understand the cortical mechanisms of auditory selective attention. Hearing Research, 307, 111–120. 10.1016/J.HEARES.2013.06.010

Lesenfants, D., Vanthornhout, J., Verschueren, E., Decruy, L., & Francart, T. (2019). Predicting individual speech intelligibility from the cortical tracking of acoustic- and phonetic-level speech representations. Hearing Research, 380, 1–9. 10.1016/J.HEARES.2019.05.006

Liberman, M. C. (1982). The cochlear frequency map for the cat: Labeling auditory-nerve fibers of known characteristic frequency. The Journal of the Acoustical Society of America, 72(5), 1441–1449. 10.1121/1.388677

Liu, T. C., Lin, Y. C., Wang, T. N., Yeh, S. C., & Kalyuga, S. (2021). Studying the effect of redundancy in a virtual reality classroom. Educational Technology Research and Development, 69(2), 1183–1200. 10.1007/S11423-021-09991-6/TABLES/1

Lozano-Soldevilla, D. (2018). On the physiological modulation and potential mechanisms underlying parieto-occipital alpha oscillations. Frontiers in Computational Neuroscience, 12, 340698. 10.3389/FNCOM.2018.00023/BIBTEX

Maamor, N., & Billings, C. J. (2017). Cortical signal-in-noise coding varies by noise type, signal-to-noise ratio, age, and hearing status. Neuroscience Letters, 636, 258–264. 10.1016/J.NEULET.2016.11.020

Mackersie, C. L., & Cones, H. (2011). Subjective and psychophysiological indexes of listening effort in a competing-talker task. Journal of the American Academy of Audiology, 22(2), 113–122. 10.3766/JAAA.22.2.6

Mackersie, C. L., Macphee, I. X., & Heldt, E. W. (2015). Effects of Hearing Loss on Heart-Rate Variability and Skin Conductance Measured During Sentence Recognition in Noise. Ear and Hearing, 36(1), 145. 10.1097/AUD.0000000000000091

Makov, S., Sharon, O., Ding, N., Ben-Shachar, M., Nir, Y., & Golumbic, E. Z. (2017). Sleep Disrupts High-Level Speech Parsing Despite Significant Basic Auditory Processing. Journal of Neuroscience, 37(32), 7772–7781. 10.1523/JNEUROSCI.0168-17.2017

Marrufo-Pérez, M. I., Lopez-Poveda, E. A., & Marrufo-P Erez, M. I. (2022). Adaptation to noise in normal and impaired hearing. The Journal of the Acoustical Society of America, 151(3), 1741–1753. 10.1121/10.0009802

Martin, D., & Miller, C. (2012). Speech and Language Difficulties in the Classroom. Speech and Language Difficulties in the Classroom. 10.4324/9780203421109

Mattys, S. L., Davis, M. H., Bradlow, A. R., & Scott, S. K. (2012). Speech recognition in adverse conditions: A review. Language and Cognitive Processes, 27(7–8), 953–978. 10.1080/01690965.2012.705006

Miles, K., McMahon, C., Boisvert, I., Ibrahim, R., de Lissa, P., Graham, P., & Lyxell, B. (2017a). Objective Assessment of Listening Effort: Coregistration of Pupillometry and EEG. Trends in Hearing, 21. 10.1177/2331216517706396

Miles, K., McMahon, C., Boisvert, I., Ibrahim, R., de Lissa, P., Graham, P., & Lyxell, B. (2017b). Objective Assessment of Listening Effort: Coregistration of Pupillometry and EEG. Trends in Hearing, 21. 10.1177/2331216517706396/ASSET/IMAGES/LARGE/10.1177_2331216517706396-FIG4.JPEG

Miller, G. A., & Licklider, J. C. R. (1950). The Intelligibility of Interrupted Speech. The Journal of the Acoustical Society of America, 22(2), 167–173. 10.1121/1.1906584

Mueller-Pfeiffer, C., Zeffiro, T., O’Gorman, R., Michels, L., Baumann, P., Wood, N., Spring, J., Rufer, M., Pitman, R. K., & Orr, S. P. (2014). Cortical and cerebellar modulation of autonomic responses to loud sounds. Psychophysiology, 51(1), 60–69. 10.1111/PSYP.12142

Niemczak, C. E., & Vander Werff, K. R. (2019). Informational Masking Effects on Neural Encoding of Stimulus Onset and Acoustic Change. Ear and Hearing, 40(1), 156–167. 10.1097/AUD.0000000000000604

Obleser, J., Wöstmann, M., Hellbernd, N., Wilsch, A., & Maess, B. (2012). Adverse Listening Conditions and Memory Load Drive a Common Alpha Oscillatory Network. Journal of Neuroscience, 32(36), 12376–12383. 10.1523/JNEUROSCI.4908-11.2012

Orf, M., Wöstmann, M., Hannemann, R., & Obleser, J. (2023). Target enhancement but not distractor suppression in auditory neural tracking during continuous speech. IScience, 26(6), 106849. 10.1016/J.ISCI.2023.106849

O’Sullivan, J. A., Power, A. J., Mesgarani, N., Rajaram, S., Foxe, J. J., Shinn-Cunningham, B. G., Slaney, M., Shamma, S. A., & Lalor, E. C. (2015). Attentional Selection in a Cocktail Party Environment Can Be Decoded from Single-Trial EEG. Cerebral Cortex, 25(7), 1697–1706. 10.1093/CERCOR/BHT355

Palva, S., & Palva, J. M. (2011). Functional Roles of Alpha-Band Phase Synchronization in Local and Large-Scale Cortical Networks. Frontiers in Psychology, 2(SEP). 10.3389/FPSYG.2011.00204

Parsons, T. D. (2015). Virtual reality for enhanced ecological validity and experimental control in the clinical, affective and social neurosciences. Frontiers in Human Neuroscience, 9(DEC), 146520. 10.3389/FNHUM.2015.00660/BIBTEX

Peelle, J. E. (2018). Listening Effort: How the Cognitive Consequences of Acoustic Challenge Are Reflected in Brain and Behavior. Ear and Hearing, 39(2), 204. 10.1097/AUD.0000000000000494

Rimmele, J. M., Zion Golumbic, E., Schröger, E., & Poeppel, D. (2015). The effects of selective attention and speech acoustics on neural speech-tracking in a multi-talker scene. Cortex, 68, 144–154. 10.1016/J.CORTEX.2014.12.014

Risko, E. F., Richardson, D. C., & Kingstone, A. (2016). Breaking the Fourth Wall of Cognitive Science: Real-World Social Attention and the Dual Function of Gaze. Current Directions in Psychological Science, 25(1), 70–74. 10.1177/0963721415617806

Šabić, E., Henning, D., Myüz, H., Morrow, A., Hout, M. C., & MacDonald, J. A. (2020). Examining the Role of Eye Movements During Conversational Listening in Noise. Frontiers in Psychology, 11, 500343. 10.3389/FPSYG.2020.00200/BIBTEX

Sala, E., & Rantala, L. (2016). Acoustics and activity noise in school classrooms in Finland. Applied Acoustics, 114, 252–259. 10.1016/J.APACOUST.2016.08.009

Salmelin, R. (2007). Clinical neurophysiology of language: The MEG approach. Clinical Neurophysiology, 118(2), 237–254. 10.1016/J.CLINPH.2006.07.316

Schneider, D., Herbst, S. K., Klatt, L. I., & Wöstmann, M. (2022). Target enhancement or distractor suppression? Functionally distinct alpha oscillations form the basis of attention. European Journal of Neuroscience, 55(11–12), 3256–3265. 10.1111/EJN.15309

Shavit-Cohen, K., & Zion Golumbic, E. (2019). The Dynamics of Attention Shifts Among Concurrent Speech in a Naturalistic Multi-speaker Virtual Environment. Frontiers in Human Neuroscience, 13, 469813. 10.3389/FNHUM.2019.00386/BIBTEX

Shield, B., Conetta, R., Dockrell, J., Connolly, D., Cox, T., & Mydlarz, C. (2015). A survey of acoustic conditions and noise levels in secondary school classrooms in England. The Journal of the Acoustical Society of America, 137(1), 177–188. 10.1121/1.4904528

Shield, B., & Dockrell, J. E. (2004). External and internal noise surveys of London primary schools. The Journal of the Acoustical Society of America, 115(2), 730–738. 10.1121/1.1635837

Shinn-Cunningham, B. G., & Best, V. (2008). Selective attention in normal and impaired hearing. Trends in Amplification, 12(4), 283–299. 10.1177/1084713808325306

Song, J., & Iverson, P. (2018). Listening effort during speech perception enhances auditory and lexical processing for non-native listeners and accents. Cognition, 179, 163–170. 10.1016/J.COGNITION.2018.06.001

Strauß, A., Kotz, S. A., & Obleser, J. (2013). Narrowed Expectancies under Degraded Speech: Revisiting the N400. Journal of Cognitive Neuroscience, 25(8), 1383–1395. 10.1162/JOCN_A_00389

Strauß, A., Wöstmann, M., & Obleser, J. (2014). Cortical alpha oscillations as a tool for auditory selective inhibition. Frontiers in Human Neuroscience, 8(MAY), 88192. 10.3389/FNHUM.2014.00350/BIBTEX

Ten Oever, S., Schroeder, C. E., Poeppel, D., Van Atteveldt, N., Mehta, A. D., Mégevand, P., Groppe, D. M., & Zion-Golumbic, E. (2017). Low-Frequency Cortical Oscillations Entrain to Subthreshold Rhythmic Auditory Stimuli. The Journal of Neuroscience_: The Official Journal of the Society for Neuroscience, 37(19), 4903–4912. 10.1523/JNEUROSCI.3658-16.2017

Thompson, R. F., & Spencer, W. A. (1966). Habituation: a model phenomenon for the study of neuronal substrates of behavior. Psychological Review, 73(1), 16–43. 10.1037/H0022681

Tromp, J., Peeters, D., Meyer, A. S., & Hagoort, P. (2018). The combined use of virtual reality and EEG to study language processing in naturalistic environments. Behavior Research Methods, 50(2), 862–869. 10.3758/S13428-017-0911-9/FIGURES/3

Uther, M., Jansen, D. H. J., Huotilainen, M., Ilmoniemi, R. J., & Näätänen, R. (2003). Mismatch negativity indexes auditory temporal resolution: evidence from event-related potential (ERP) and event-related field (ERF) recordings. Cognitive Brain Research, 17(3), 685–691. 10.1016/S0926-6410(03)00194-0

Vanthornhout, J., Decruy, L., Wouters, J., Simon, J. Z., & Francart, T. (2018). Speech Intelligibility Predicted from Neural Entrainment of the Speech Envelope. JARO - Journal of the Association for Research in Otolaryngology, 19(2), 181–191. 10.1007/S10162-018-0654-Z/FIGURES/6

Verschueren, E., Vanthornhout, J., & Francart, T. (2020). The Effect of Stimulus Choice on an EEG-Based Objective Measure of Speech Intelligibility. Ear and Hearing, 41(6), 1586–1597. 10.1097/AUD.0000000000000875

Wöstmann, M., Herrmann, B., Wilsch, A., & Obleser, J. (2015). Neural Alpha Dynamics in Younger and Older Listeners Reflect Acoustic Challenges and Predictive Benefits. Journal of Neuroscience, 35(4), 1458–1467. 10.1523/JNEUROSCI.3250-14.2015

Yasmin, S., Irsik, V. C., Johnsrude, I. S., & Herrmann, B. (2023). The effects of speech masking on neural tracking of acoustic and semantic features of natural speech. Neuropsychologia, 186, 108584. 10.1016/J.NEUROPSYCHOLOGIA.2023.108584

Zion Golumbic, E. M., Ding, N., Bickel, S., Lakatos, P., Schevon, C. A., McKhann, G. M., Goodman, R. R., Emerson, R., Mehta, A. D., Simon, J. Z., Poeppel, D., & Schroeder, C. E. (2013). Mechanisms Underlying Selective Neuronal Tracking of Attended Speech at a “Cocktail Party.” Neuron, 77(5), 980–991. 10.1016/J.NEURON.2012.12.037

Zou, J., Feng, J., Xu, T., Jin, P., Luo, C., Zhang, J., Pan, X., Chen, F., Zheng, J., & Ding, N. (2019). Auditory and language contributions to neural encoding of speech features in noisy environments. NeuroImage, 192, 66–75. 10.1016/J.NEUROIMAGE.2019.02.047

